# Mineralocorticoid Receptor Antagonism Reduces Atrial Arrhythmias Post-Cardiac Surgery and Attenuates Atrial Stress Responses to Cardioplegic Arrest

**DOI:** 10.1101/2025.02.24.639999

**Authors:** Sina Danesh, Fazal Khan, Trevor Chopko, Aurora Lee, Ran Huo, Shuyang Lu, Vincy Tam, Pedro Fincatto Safi, Wenbin Gao, Austin Todd, Francis D. Pagani, Joseph J. Maleszewski, Hartzell Schaff, Paul A. Friedman, Hakan Oral, Marco Metra, Bertram Pitt, Ienglam Lei, Paul C. Tang

## Abstract

**Background:** New postoperative atrial fibrillation (POAF) occurs in about 40% after cardiac surgery. Mineralocorticoid receptor antagonists (MRA) are known to reduce chronic atrial fibrillation (AF) development and burden. We examined the impact of preoperative MRA use on POAF and also examine the atrial cell type impacted by MRA treatment during cold cardiac preservation.

**Methods:** Retrospective study of 19,042 patients who underwent cardiac surgery at Mayo Clinic in Minnesota, and performed 1:3 propensity matching to obtain 298 patients on preoperative MRA matched to 894 who were not. We also separately matched patients using preoperative diuretics. Single-nuclei RNA sequencing (snRNA-seq) examined MRA’s effects on different atrial cell types in canrenone (water soluble MRA) treated human donor hearts undergoing cold preservation followed by ex-vivo reperfusion and compared gene expression to the atria of patients with AF.

**Results:** Propensity matched preoperative MRA group had less new onset POAF (19.8% vs 31.5%, P<0.001). To account for the possibility that preoperative diuretic use and volume reduction may impact POAF, we propensity matched 298 preop diuretic users that included MRA use to another 894 patients who used a non-MRA diuretic preoperatively. Those who used preoperative MRA similarly had a lower incidence of POAF (19.8% vs 33.2%, P<0.001). No survival difference was present between the propensity matched groups that used preoperative diuretics (P=0.079). Preoperative MRA use also reduced the development of paroxysmal and chronic AF at 6 years of follow up. From our snRNA-seq data, we identified a subpopulation of atrial cardiomyocytes (CM2) that had high MR expression where canrenone suppressed the increase in MR target gene expression associated with cold preservation-reperfusion. These MR targets were conversely elevated in patients with chronic AF. Canrenone also suppressed other cardiac preservation associated genes that show elevated expression in atrial macrophages and pericytes from chronic AF atria.

**Conclusions:** Our studies show that preoperative MRA use is associated with 40% reduction in POAF as well as lowering long standing AF development by about 41%. Our cold cadiac preservation-reperfusion model showed that canrenone reduced expression of MR target genes associated with chronic AF, particular in cardiomyocytes with important roles in electrical conduction.

**Clinical Perspective:** *What is New?:*

- This study shows that preoperative use of mineralocorticoid receptors antagonists (MRA) is associated with a reduced incidence of new onset perioperative atrial fibrillation after cardiac surgery utilizing cardiopulmonary bypass.
- We show that preoperative MRA use is associated with a lower incidence of developing more chronic paroxymal or sustained atrial fibrillation.
- Addition of canrenone, a clinically utilized water soluble MRA, to cardioplegia solution used during cardiac preservation can attenuate atrial inflammatory reponses and reduce signaling through molecular pathways that promote atrial fibrillation.

*What are the clinical implications?:*

- Perioperative use of MRAs may be considered to reduce early postoperative atrial fibrillation as well as lowering the risk of developing more chronic atrial arrhythmias.
- These findings support pursuing a clinical trial to determine the impact of MRA use on atrial arrhythmias following cardiac surgery in the setting of cardiopulmonary bypass with cold cardiac preservation.

## INTRODUCTION

New onset postoperative atrial fibrillation (POAF) after cardiac surgery is associated with increased perioperative complications and early mortality^1–3^. POAF incidence range from 20-40% after coronary artery bypass grafting with or without aortic valve replacement and 64% after combined CABG and mitral surgery. POAF rates are even higher in patients with heart failure^4–6^. Importantly, patients with new POAF after cardiac surgery have a high incidence of late atrial fibrillation (AF) recurrence of 10-49% over 3.4-8.3 years^7–10^. Thus, acute POAF episodes begets more long term occurrence of AF with implications extending far beyond the postoperative setting. POAF is recognized as multifactorial with contibutions from underlying co-morbidities, operative stress, cardiac preservation and reperfusion injury during cardioplegic arrest, fluctuations in intra-cardiac pressures as well as intrinsic and extrinsic hormonal exposures (e.g. catecholamines)^4,11–13^.

POAF incidence remains high despite the wide use of β-blockers and amiodarone to prevent and treat this complication^14^. Mineralocorticoid receptor antagonists (MRA, e.g. spironolactone, eplerenone, finerenone) are well recognized to reduce AF burden in various nonsurgical settings such as heart failure and diabetes^15–19^. The role of mineralocorticoid receptor (MR) in POAF is remains unclear although individuals with hyperaldosteronism are known to have a higher rate of cardiovascular events including a 12-fold higher incidence of AF that is independent of hypertension^20^. This is attributed to MR signaling activating the substrates for AF such as atrial fibrosis, modulation of ion channel expression as well as myocardial inflammation and oxidative stress^21–24^. Interestingly, patients with chronic AF have elevated baseline serum aldosterone levels but these decline after reverting to sinus rhythm^25^.

Given these findings, we examined the effects of preoperative MRA use and show that it is associated with a much lower incidence of POAF after cardiac surgery. Importantly, we show that preoperative MRA reduces the development of AF in the longer term. We also gained insights through translational science studies on the impact of MRAs on cellular signaling in human left atrial tissue at the single-nuclei level. We anticipate these results to be clinically relevant for developing peri- and intraoperative targeted therapies to reduce POAF and importantly, lower the lifetime risk of AF which may be triggered by cardiac surgery.

## METHODS

### Clinical Study Population

Our clinical study design with a waiver of informed consent was approved by the Mayo Clinic Institutional Review Board (IRB, 23-009355, approved October 10^th^, 2023). We conducted a retrospective review of 45,368 patients who underwent open cardiac surgery using cardiopulmonary bypass from January 5^th^, 1993, to December 29^th^, 2023 at the Mayo Clinic Department of Cardiovascular Surgery in Rochester, Minnesota. The study sample included adults aged ≥18 years and less than 75 year of age undergoing a “primary sternotomy” for isolated or combined surgeries that included coronary artery bypass grafting, myectomy for hypertropic cardiomyopathy, and/or valve surgeries (aortic, mitral, tricuspid and/or pulmonary valve). We excluded patients with pre-existing atrial arrhythmias (atrial fibrillation or flutter), prior cardiac surgery using cardiopulmonary bypass, cardiac surgery for emergent indications, or surgeries not mentioned previously (e.g. aortic procedures, pericardiotomy, pulmonary thromboenarterectomy, thoracic transplantation, ventricular assist device implants). Patients who underwent concomitant atrial MAZE procedure or left atrial appendage excision or ligation were also excluded from the study. The final study cohort consisted of 19,042 patients of which 319 patients were on MRAs preoperatively. Graphical representation of the inclusion and exclusion criteria and development of the study cohort is presented in **Figure 1**.

**Figure 1.**
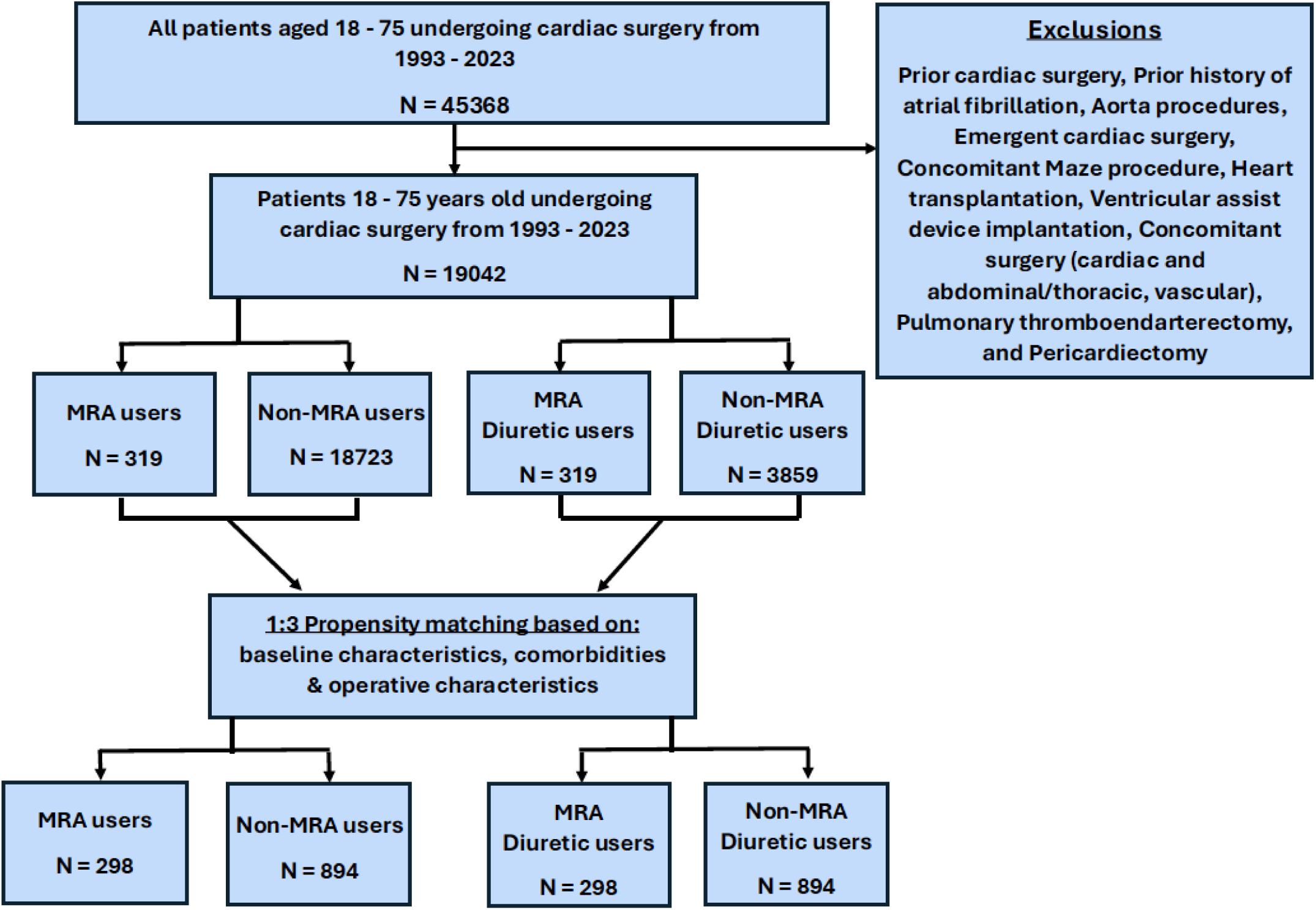
Consort diagram detailing the application of inclusion and exclusion criteria.

### Clinical Study Endpoints

We examined patient demographics, comorbidities, preoperative medication use, intraoperative parameters and postoperative occurrence of atrial arrhythmias. Postoperative atrial arrhythmias was defined as per Society of Thoracic Surgeons (STS) criteria as AF that occurs within 30 days after surgery and requires treatment or lasts longer than one hour^26^. The primary endpoint of interest in the study was the occurrence of postoperative new onset AF. Secondary endpoints includes patient survival, postoperative infection, postcardiotomy shock, renal failure, cerebrovascular complications, gatrointestinal bleeding, blood product use, permanent pacemaker use and length of hospital stay.

### Propensity Matching

Covariates were analyzed with logistic regression to generate a propensity score for propensity toward being on a MRA prior to undergoing cardiac surgery. We performed a 1:3 greedy matching algorithm without replacement using a caliper width of 0.02 standard deviations of the linear predictor (Figure 1). Matched variables included patient age, sex, body mass index, creatinine, hypertension, diabetes, left ventricular ejection fraction (LVEF), preoperative β-blocker and amiodarone use as well as history of transient ischemic attacks, cerebrovascular accidents, myocardial infarction and chronic lung disease. We also matched for intraoperative parameters including cardiopulmmonary bypass time and aortic cross-clamp time as well as performance of coronary artery bypass grafting, myectomy for hypertrophic cardiomyopathy and valve surgeries (aortic, mitral, tricuspid, pulmonic valve repair or replacement). Following matching, there were 298 patients in the MRA group and 894 patients in the no MRA group. The reduction of atrial stretch and volume overload and electrolyte changes from preoperative diuretic use may impact the burden of postoperative atrial arrhythmias^27–31^. To separate the effects of diuretics drugs on volume and electrolyte status from cardiac specific effects of MRAs, we performed a subanalysis on patients using diuretics preoperatively and similarly performed propensity matching for preoperative MRA use (n=298) or not (n=894)

### Data collection

A prospectively maintained institutional cardiovascular surgery database was used to obtain relevant demographic data, comorbidities, echocardiographic data, operative details, as well as postoperative and mid-term outcomes. The variables were defined on the basis of the criteria set forth in the Society of Thoracic Surgeons (STS) Adult Cardiac Surgery Database (STS-ACSD). We conducted a 6-year follow-up using medical records to evaluate the severity of POAF in patients who developed the condition. Clinical details not included in the database such as development of chronic atrial arrhythmias, medication dosage and specific drug classes were extracted through retrospective chart review.

### Statistical Analysis

Categorical data are presented as frequency (percentage of group), with comparisons using Pearson’s Chi-squared test. Continuous variables are expressed as the median (25^th^-75^th^ percentile), with categorical variables displayed as numbers (percentages). Wilcoxon rank sum test was used to compare demographics, baseline characteristics, and operative parameters between patients. Patient survival was analyzed by Kaplan-Meir methodology with log-rank statistics. The relationship between preoperative MRA use with POAF and survival were also assessed using a multivariable Cox proportional hazard model. Analyses were conducted using R software version 4.1.3 with p-values < 0.05 considered statistically significant.

### Research ethics for organ donor and surgical heart tissues

Human donor heart studies were approved by the University of Michigan Institutional Review Board (HUM00131275) and Mayo Clinic Institutional Review Board (23-006893). Atrial specimens were obtained from patients undergoing heart surgery after securing informed consent. Mayo Clinic Institutional Review Board approval (23-008400, 23-007374) was also obtained for surgical tissue.

### Procurement and preparation of human donor atrial tissue

Heart procurement for humans were performed as per clinical standard. A sternotomy is performed under general anesthesia. The pericardium is then opened to create a pericardial cradle for the heart. Heparin (300U/kg) is administered in a peripheral vein (for humans). After waiting 3 minutes for adequate anticoagulation from heparin, a 9 Fr antegrade plegia infusion catheter is placed in the ascending aorta. Approximately 1 liter of autologous donor arterial blood is then collected into a collection bottle containing heparin. The human or pig donor heart is then retrieved per clinical protocol by incising the inferior vena cava to drain the right heart and then the left atrial appendage is incised to drain the left heart. Subsequently, the ascending aorta is cross-clamped distal to the cardioplegia catheter and 1 liter of cold (4°C) HTK solution is infused into the coronaries at a perfusion pressure of ∼80 mmHg to induce mechanical arrest. After excising the heart completely, it is transported to the back table for infusion of 2 liters of HTK±Canrenone (50 μM) solution. After completion, the heart is stored in a solution identical to the perfusate at 4°C on ice. Human hearts underwent ex-vivo perfusion after cold (4°C) static preservation for 10 hours. The left atrial appendage was collected and were either fixed in formalin or snap frozen in liquid nitrogen for later analysis. Clinical details such as demographics and comorbidities of human heart donors as well as cardiac surgery patients donating left atrial appendages are provided in “Supplementary Data”.

### Human ex-vivo heart perfusion

Details of large animal heart perfusion were described previously^32^. Briefly, the ascending aorta is accessed using an arterial perfusion cannula and the pulmonary veins are sutured shut to close the left atrium. A catheter is placed in the left ventricle through the left atrium as a vent. The Radnoti constant perfusion pressure apparatus (ADInstruments Inc., Colorado Springs, CO) was utilized for these human ex-vivo heart perfusion studies. About 1 liter of autologous blood was added to ∼3.5 liters of Krebs buffer. This was initially infused at 24°C with progressive warming to 37°C. Great care is taken to remove air from the perfusion system. The ascending aorta and coronaries were then retrograde perfused through the arterial cannula at ∼80 mmHg with subsequent cardiac reanimation. The perfusion pressure was kept constant at ∼80 mmHg so LV contraction would need to overcome this afterload for cardiac output. Oxygen (pO_2_) pressure was kept at approximatey 150–200 mmHg while pCO_2_ was maintained at about 35-40 mmHg. Arterial pH was kept relateively constant at 7.35 – 7.40 via buffering with 5% CO_2_ during gas exchange. The human donor heart was ex-vivo perfused for 3 hours and then arrested with 4°C HTK solution for left atrial appendage specimen collection with liquid nitrogen.

### Single nuclei RNA-seq

Human heart tissues were collected from the left atrial appendage of hearts from organ donors and following consent from hearts of patients undergoing cardiac surgery at the Mayo Clinic in Rochester, Minnesota. Collection of specimens for this study was approved by the Mayo Clinic IRB (#23-008400, approved October 3^rd^, 2023; #23-007374; #23-006893). Singleron Biotechnologies (Ann Arbor, Michigan) performed the single nuclei RNA-seq. Single nuclei were isolated from flash-frozen appendage tissue followed by GEXSCOPE^®^ library preparation followed by library sequencing at a depth of 90gb persample. The raw sequencing results were aligned to human genome GRCh 38 and aggregated with CellRanger v7.1.03^33^. The count matrix was further analyzed with Seurat v4.4.04^34^. Cells were filtered for a RNA count <25000, RNA count >1000, and <5% containing mitochondrial gene. This filtered library was normalized with the open-source R package SCTransform (github.com/ChristophH/sctransform)^35^.

The expression matrix from different samples was integrated using the RPCAIntergration function in Seurat. Uniform Manifold Approximation and Projection (UMAP) was used to visualize data^36^ for dimension reduction. FindClusters function^37^ was used for cluster detection with the default resolution (0.8). In addition to the graphical display, we used the FindMarkers function^37^ in the Seurat package to perform differential expression gene (DEG) analysis among phenotype-associated subpopulations. We then annotated the cell types of each cluster based on the highest positive fold changes and correlated them with relevant specific cell markers described in the literature^38^. Fgsea was used to identify the gene set enrichment in the subcluster of each cell type. The expression matrix for individual cell type was extracted for further analysis. We identified the canrenone-regulated genes using Findmarkers. The expression of canrenone-regulated genes in each cell type was then estimated using the AddModuleScore function and tested with Wilcox ranking comparison.

## RESULTS

### Patient Characteristics

The study cohort included 19,042 patients who met inclusion criteria (Supplementary Table 1). Compared to preoperative non-MRA users, MRA users (n=319) have a higher incidence of females (43.6% vs 31.4%, P<0.001), hypertension (73.0% vs 62.1%, P<0.001), diabetes (35.4% vs 21.6%, P<0.001), chronic lung disease (6.6% vs 2.4%, P<0.001), preoperative β-blocker use (66.1% vs 52.5%, P<0.001), preopeartive amiodarone use (2.5% vs 0.7%, P<0.001) as well as higher Society of Thoracic Surgeons (STS) surgical mortality risk (1.4% vs 1.0%, P<0.001). MRA users also had lower GFR (61.3 vs 67.5 ml/min/1.73m^2^, P<0.001).

Echocardiography showed that MRA users has lower left ventricular ejection fraction (LVEF, 59.5% vs 63.0%, P<0.001) and a higher incidence of valve pathology including significant aortic regurgitation (15.6% vs 10.8%, P=0.005), aortic stenosis (27.9% vs 22.3%, P=0.017), mitral regurgitation (35.4% vs 33.5%, P<0.001), mitral stenosis (9.1% vs 3.7%, P<0.001), tricuspid regurgitation (22.6% vs 6.8%, P<0.001) and pulmonary regurgitation (8.1% vs 1.8%, P<0.001). Accordingly, MRA users had more valve procedures including aortic valve replacement (29.8% vs 23.3%, P=0.007), mitral valve replacement (8.8% vs 4.0%, P<0.001), tricuspid valve replacement (8.2% vs 1.2%, P<0.001), tricuspid valve repair (6.3% vs 1.9%, P<0.001), and pulmonic valve replacement (5.3% vs 0.8%, P<0.001). However, MRA users had less mitral valve repair (16.3% vs 21.6%, P=0.023). MRA users also had higher cardiopulmonary bypass (83.0 min vs 76.0 min, P<0.001) and aortic cross clamp times (59.0 min vs 52.0 min, P<0.001).

### Propensity matched groups and new onset atrial arrhythmias

Following 1:3 matching of the entire study population, there were 298 patients in the MRA user group and 894 patients in the non-MRA group (Supplementary Table 2). Propensiy score distribution for the entire study population as well as jitter plot and histogram for propensity score distribution in the matched cohorts are shown in Supplementary Figures 1 and 2. POAF incidence was lower in the MRA user group (19.8% vs 31.5%, P<0.001) compared to non-MRA users. There were no differences in other identified outcome measures (Supplementary Table 3).

Volume management using diuretics may impact atrial stretch and the development of atrial arrhytmias^27–31^. In an effort to focus on cardiac specific effects of MRAs beyond renal diuresis, we subsequently limited our analysis to patients on preoperative diuretics (n=4,178) where clinical demographics were more comparable between MRA (n=319) and non-MRA diuretic users (n=3,859). However, MRA users still had a greater burden of valvular pathologies and related procedures (Supplementary Table 4). We again performed propensity matching to obtain comparable groups of MRA (n=298) and non-MRA (n=894) users (Table 1). Propensiy score distribution for the entire study population as well as jitter plot and histogram for propensity score distribution in the matched cohorts are shown in Supplementary Figures 3 and 4.

**Table 1.**
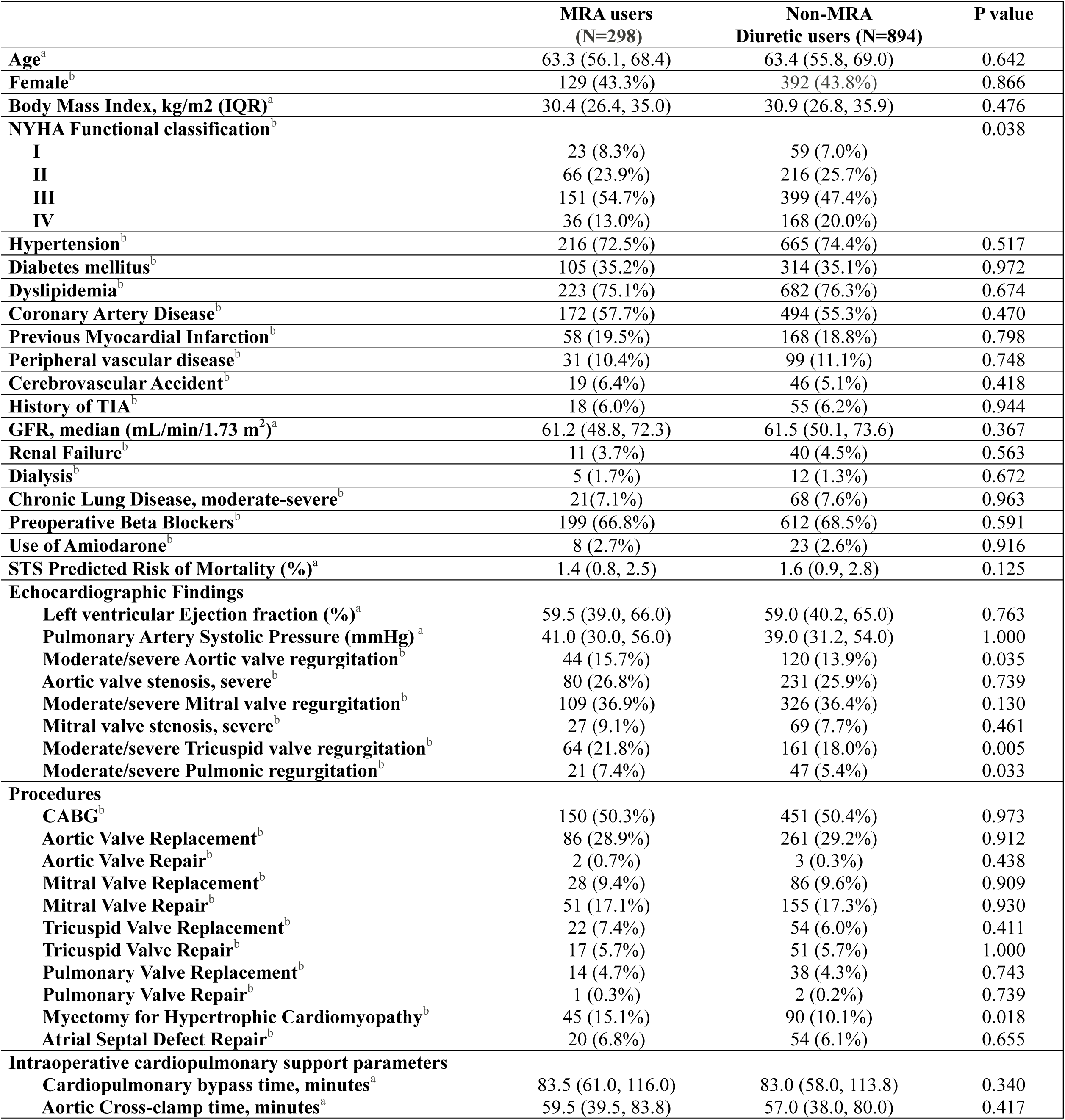
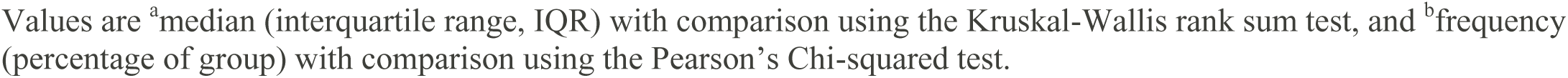
Baseline Demographics, Pathology, and Operative Characteristics of Diuretic Users Stratified by Using Mineralocorticoid Receptor Antagonist (MRA) After Propensity Matching.

|  | **MRA users**<br>**(N=298)** | **Non-MRA**<br>**Diuretic users (N=894)** | **P value** |
| --- | --- | --- | --- |
| **Age<sup>a</sup>** | 63.3 (56.1, 68.4) | 63.4 (55.8, 69.0) | 0.642 |
| **Female<sup>b</sup>** | 129 (43.3%) | 392 (43.8%) | 0.866 |
| **Body Mass Index, kg/m2 (IQR)<sup>a</sup>** | 30.4 (26.4, 35.0) | 30.9 (26.8, 35.9) | 0.476 |
| **NYHA Functional classification<sup>b</sup>** |  |  | 0.038 |
| **I** | 23 (8.3%) | 59 (7.0%) |  |
| **II** | 66 (23.9%) | 216 (25.7%) |  |
| **III** | 151 (54.7%) | 399 (47.4%) |  |
| **IV** | 36 (13.0%) | 168 (20.0%) |  |
| **Hypertension<sup>b</sup>** | 216 (72.5%) | 665 (74.4%) | 0.517 |
| **Diabetes mellitus<sup>b</sup>** | 105 (35.2%) | 314 (35.1%) | 0.972 |
| **Dyslipidemia<sup>b</sup>** | 223 (75.1%) | 682 (76.3%) | 0.674 |
| **Coronary Artery Disease<sup>b</sup>** | 172 (57.7%) | 494 (55.3%) | 0.470 |
| **Previous Myocardial Infarction<sup>b</sup>** | 58 (19.5%) | 168 (18.8%) | 0.798 |
| **Peripheral vascular disease<sup>b</sup>** | 31 (10.4%) | 99 (11.1%) | 0.748 |
| **Cerebrovascular Accident<sup>b</sup>** | 19 (6.4%) | 46 (5.1%) | 0.418 |
| **History of TIA<sup>b</sup>** | 18 (6.0%) | 55 (6.2%) | 0.944 |
| **GFR, median (mL/min/1.73 m<sup>2</sup>)<sup>a</sup>** | 61.2 (48.8, 72.3) | 61.5 (50.1, 73.6) | 0.367 |
| **Renal Failure<sup>b</sup>** | 11 (3.7%) | 40 (4.5%) | 0.563 |
| **Dialysis<sup>b</sup>** | 5 (1.7%) | 12 (1.3%) | 0.672 |
| **Chronic Lung Disease, moderate-severe<sup>b</sup>** | 21 (7.1%) | 68 (7.6%) | 0.963 |
| **Preoperative Beta Blockers<sup>b</sup>** | 199 (66.8%) | 612 (68.5%) | 0.591 |
| **Use of Amiodarone<sup>b</sup>** | 8 (2.7%) | 23 (2.6%) | 0.916 |
| **STS Predicted Risk of Mortality (%)<sup>a</sup>** | 1.4 (0.8, 2.5) | 1.6 (0.9, 2.8) | 0.125 |
| **Echocardiographic Findings** |  |  |  |
| **Left ventricular Ejection fraction (%)<sup>a</sup>** | 59.5 (39.0, 66.0) | 59.0 (40.2, 65.0) | 0.763 |
| **Pulmonary Artery Systolic Pressure (mmHg)<sup>a</sup>** | 41.0 (30.0, 56.0) | 39.0 (31.2, 54.0) | 1.000 |
| **Moderate/severe Aortic valve regurgitation<sup>b</sup>** | 44 (15.7%) | 120 (13.9%) | 0.035 |
| **Aortic valve stenosis, severe<sup>b</sup>** | 80 (26.8%) | 231 (25.9%) | 0.739 |
| **Moderate/severe Mitral valve regurgitation<sup>b</sup>** | 109 (36.9%) | 326 (36.4%) | 0.130 |
| **Mitral valve stenosis, severe<sup>b</sup>** | 27 (9.1%) | 69 (7.7%) | 0.461 |
| **Moderate/severe Tricuspid valve regurgitation<sup>b</sup>** | 64 (21.8%) | 161 (18.0%) | 0.005 |
| **Moderate/severe Pulmonic regurgitation<sup>b</sup>** | 21 (7.4%) | 47 (5.4%) | 0.033 |
| **Procedures** |  |  |  |
| **CABG<sup>b</sup>** | 150 (50.3%) | 451 (50.4%) | 0.973 |
| **Aortic Valve Replacement<sup>b</sup>** | 86 (28.9%) | 261 (29.2%) | 0.912 |
| **Aortic Valve Repair<sup>b</sup>** | 2 (0.7%) | 3 (0.3%) | 0.438 |
| **Mitral Valve Replacement<sup>b</sup>** | 28 (9.4%) | 86 (9.6%) | 0.909 |
| **Mitral Valve Repair<sup>b</sup>** | 51 (17.1%) | 155 (17.3%) | 0.930 |
| **Tricuspid Valve Replacement<sup>b</sup>** | 22 (7.4%) | 54 (6.0%) | 0.411 |
| **Tricuspid Valve Repair<sup>b</sup>** | 17 (5.7%) | 51 (5.7%) | 1.000 |
| **Pulmonary Valve Replacement<sup>b</sup>** | 14 (4.7%) | 38 (4.3%) | 0.743 |
| **Pulmonary Valve Repair<sup>b</sup>** | 1 (0.3%) | 2 (0.2%) | 0.739 |
| **Myectomy for Hypertrophic Cardiomyopathy<sup>b</sup>** | 45 (15.1%) | 90 (10.1%) | 0.018 |
| **Atrial Septal Defect Repair<sup>b</sup>** | 20 (6.8%) | 54 (6.1%) | 0.655 |
| **Intraoperative cardiopulmonary support parameters** |  |  |  |
| **Cardiopulmonary bypass time, minutes<sup>a</sup>** | 83.5 (61.0, 116.0) | 83.0 (58.0, 113.8) | 0.340 |
| **Aortic Cross-clamp time, minutes<sup>a</sup>** | 59.5 (39.5, 83.8) | 57.0 (38.0, 80.0) | 0.417 |

Values are <sup>a</sup>median (interquartile range, IQR) with comparison using the Kruskal-Wallis rank sum test, and <sup>b</sup>frequency (percentage of group) with comparison using the Pearson's Chi-squared test.

In the MRA group of propensity matched patients using preoperative diuretics, 95% used spironolactone and 5% used eplerenone. Conversely, non-MRA group diuretic use consisted predominently of loop diuretics alone (59.7%) and Thiazide diuretics alone (29.8%, Table 2). Preoperative MRA dosages are shown in Supplementary Table 5 with 25 mg being the most common dose for both spironolactone (60.7%) and eplerenone (53.3%).

**Table 2.**
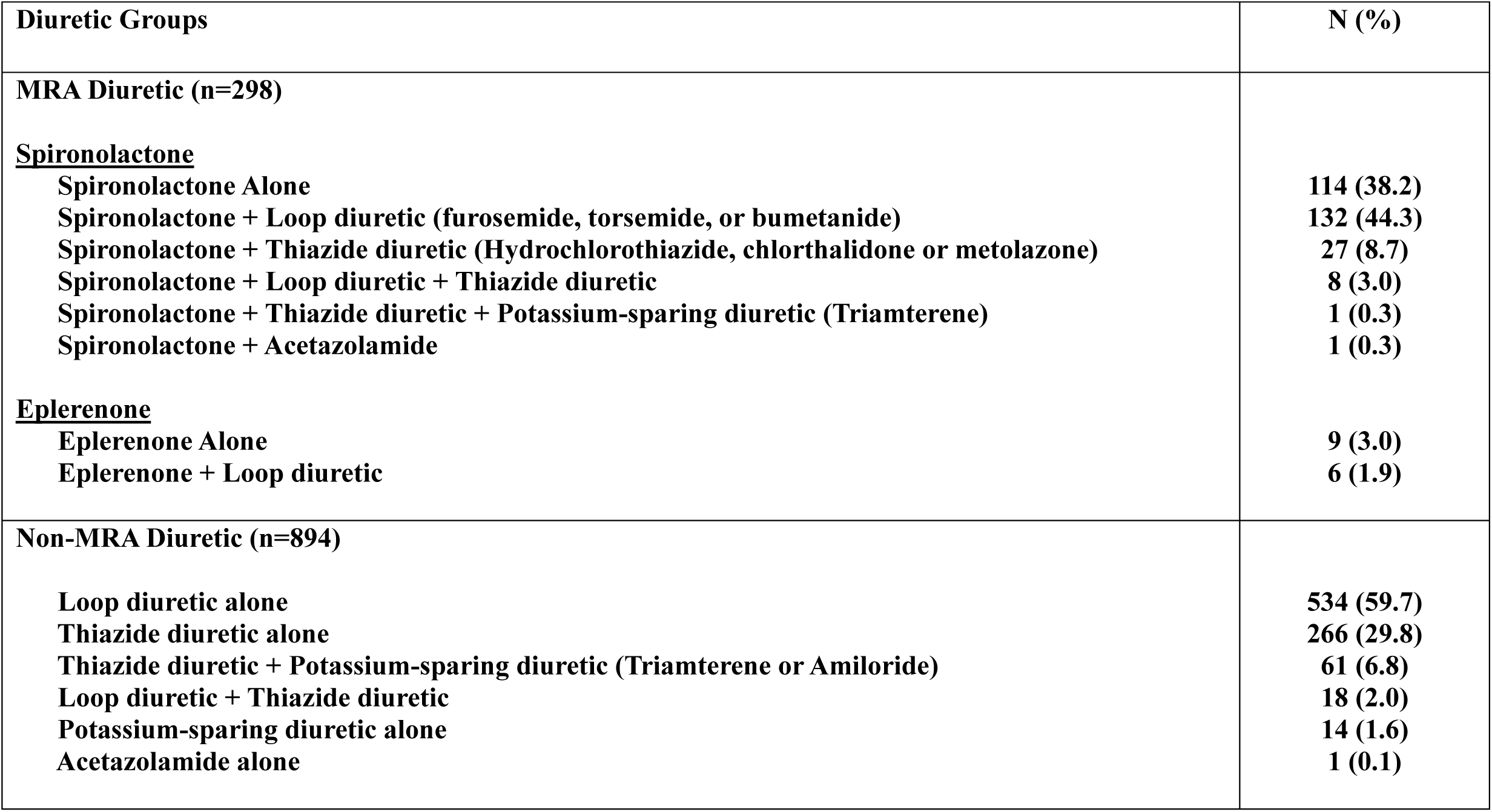
Diuretic Types in the Mineralocorticoid Receptor Antagonist (MRA) users and Non-MRA Diuretic users After Propensity Matching.

| Diuretic Groups | N (%) |
| --- | --- |
| **MRA Diuretic (n=298)** |  |
| **<u>Spironolactone</u>** |  |
| Spironolactone Alone | 114 (38.2) |
| Spironolactone + Loop diuretic (furosemide, torsemide, or bumetanide) | 132 (44.3) |
| Spironolactone + Thiazide diuretic (Hydrochlorothiazide, chlorthalidone or metolazone) | 27 (8.7) |
| Spironolactone + Loop diuretic + Thiazide diuretic | 8 (3.0) |
| Spironolactone + Thiazide diuretic + Potassium-sparing diuretic (Triamterene) | 1 (0.3) |
| Spironolactone + Acetazolamide | 1 (0.3) |
| **<u>Eplerenone</u>** |  |
| Eplerenone Alone | 9 (3.0) |
| Eplerenone + Loop diuretic | 6 (1.9) |
| **Non-MRA Diuretic (n=894)** |  |
| Loop diuretic alone | 534 (59.7) |
| Thiazide diuretic alone | 266 (29.8) |
| Thiazide diuretic + Potassium-sparing diuretic (Triamterene or Amiloride) | 61 (6.8) |
| Loop diuretic + Thiazide diuretic | 18 (2.0) |
| Potassium-sparing diuretic alone | 14 (1.6) |
| Acetazolamide alone | 1 (0.1) |

Comparing propensity matched preoperative diuretic use patients, the MRA group has a lower incidence of postoperative AF detected during their index operative admission or within 30 days of surgery at 59/298 (19.8%) versus 297/894 (33.2%, P<0.001, Table 3). At 6 years of follow up with death as a competing risk, MRA use was associated with significantly reduced the incidence of long-term combined paroxysmal or chronic atrial fibrillation (Figure 2A, P=0.003), paroxysmal AF only (Figure 2B, P=0.018) and chronic AF only (Figure 2C, P=0.081). Expressed as a categorical variable, preoperative MRA use was associated with a 40.7% reduction in long-term development of combined paroxysmal and chronic AF (38/298 (12.7%) versus 192/894 (21.4%), P=0.001), 37.1% reduction in paroxysmal AF (32/298 (10.7%) versus 152/894 (17.0%, P=0.010), and a 54.5% reduction in chronic AF (6/298 (2.0%) versus 40/894 (4.4%), P=0.056).

**Table 3.** Postoperative Outcomes and Complications of Diuretic Users Stratified by Using Mineralocorticoid Receptor Antagonist (MRA) After Propensity Matching.

| Variable | MRA users<br>(N=298) | Non-MRA<br>Diuretic users (N=894) | P value |
| --- | --- | --- | --- |
| **Postoperative Atrial Fibrillation<sup>a</sup>** | 59 (19.8%) | 297 (33.2%) | < 0.001 |
| **Readmission within 30 days<sup>a</sup>** | 26 (9.4%) | 111 (12.9%) | 0.113 |
| **Stroke<sup>a</sup>** | 5 (1.7%) | 15 (1.7%) | 1.000 |
| **TIA<sup>a</sup>** | 3 (1.1%) | 8 (0.9%) | 0.776 |
| **superficial/deep Sternal Infection<sup>a</sup>** | 6 (2.0%) | 15 (1.7%) | 0.703 |
| **Permanent Pacemaker<sup>a</sup>** | 14 (4.7%) | 56 (6.3%) | 0.319 |
| **Intra-aortic ballon pump use<sup>a</sup>** | 20 (6.7%) | 72 (8.1%) | 0.452 |
| **Extracorporeal membrane oxygenation use<sup>a</sup>** | 1 (0.3%) | 6 (0.7%) | 0.516 |
| **Cardiogenic Shock<sup>a</sup>** | 5 (1.7%) | 12 (1.3%) | 0.672 |
| **Tamponade/Pericardiocentesis<sup>a</sup>** | 6 (2.0%) | 20 (2.2%) | 0.819 |
| **Postoperative Blood Products Used<sup>b</sup>** |  |  |  |
| **Postoperative Red Blood Cells (units)<sup>c</sup>** | 2.0 (1.0, 3.0) | 2.0 (1.0, 3.0) | 0.965 |
| **Postoperative Fresh Frozen Plasma (units)<sup>c</sup>** | 2.0 (2.0, 4.0) | 2.0 (2.0, 4.0) | 0.974 |
| **Postoperative Platelet (units)<sup>c</sup>** | 1.0 (1.0, 3.0) | 1.0 (1.0, 2.0) | 0.758 |
| **Prolonged Ventilation<sup>a</sup>** | 29 (9.7%) | 91 (10.2%) | 0.824 |
| **Pneumonia<sup>a</sup>** | 8 (2.7%) | 32 (3.6%) | 0.458 |
| **Sepsis<sup>a</sup>** | 6 (2.0%) | 8 (0.9%) | 0.121 |
| **Renal failure<sup>a</sup>** | 13 (4.4%) | 39 (4.4%) | 1.000 |
| **New dialysis Required<sup>a</sup>** | 7 (2.3%) | 20 (2.2%) | 0.911 |
| **Gastrointestinal bleeding<sup>a</sup>** | 6 (2.0%) | 23 (2.6%) | 0.587 |

Values are <sup>a</sup>frequency (percentage of group) with comparison using the Pearson's Chi-squared test, <sup>b</sup>median (interquartile range, IQR) with comparison using the Kruskal-Wallis rank sum test, <sup>c</sup>frequency (percentage of group) with comparison using the Chi-squared test for given probabilities.

**Figure 2.**
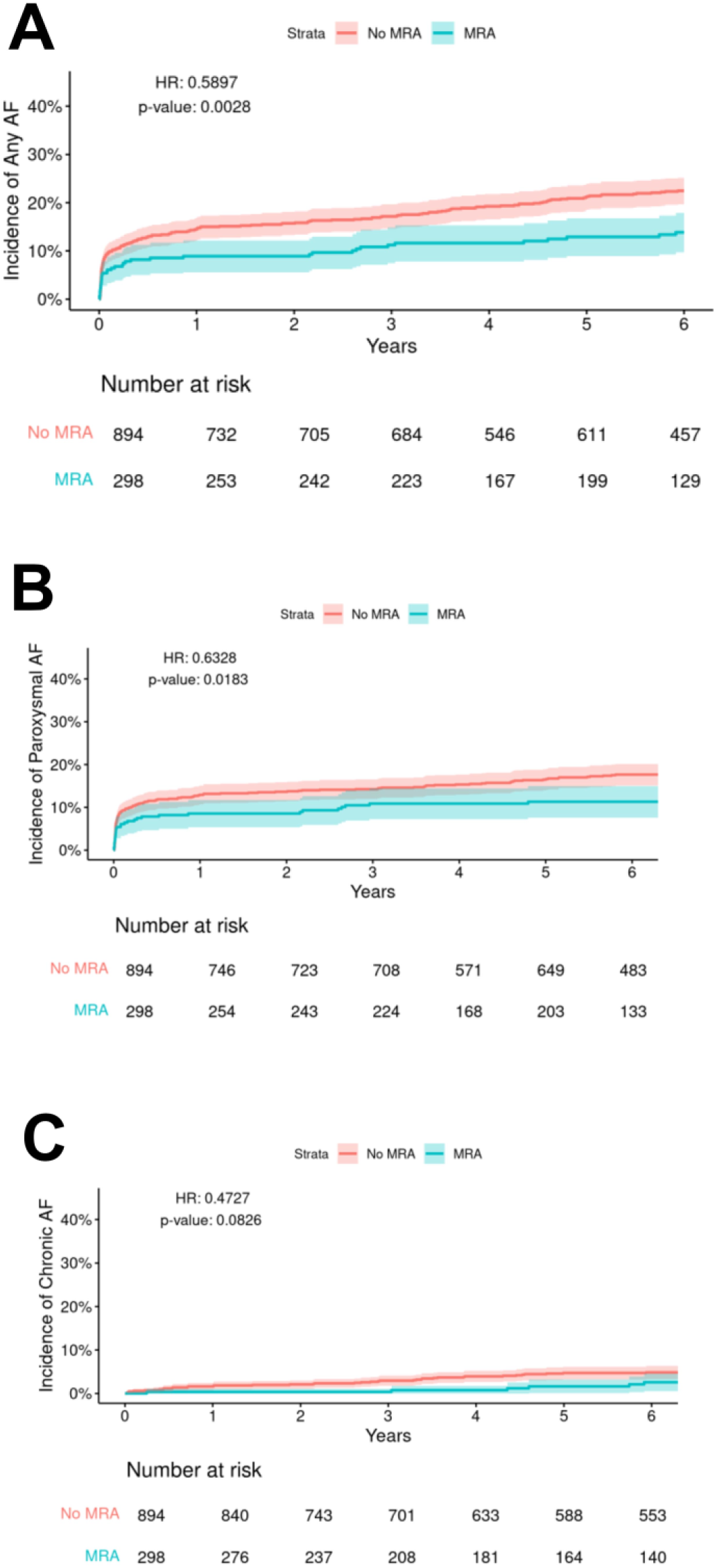
Cumulative incidence of A) Combined paroxysmal or chronic atrial fibrillation, B) Paroxysmal atrial fibrillation only and C) Chronic atrial fibrillation after cardiac surgery in propensity-matched diuretic users, stratified by preoperative mineralocorticoid receptor antagonist use with death as a competing risk (95% CI).

### Patient Survival

Preoperative MRA patients had poorer long term survival in both the total study population (OR=0.45, P<0.001) as well as the propensity matched cohort (OR=0.71, P<0.001, Supplementary Figure 5A-B). However, there was no significant difference in survival when examining only propensity matched populations who were on preoperative diuretics (OR=0.84, P=0.079, Supplementary Figure 5A-B).

### MRA suppressed the atrial expression of MR target gene sets that are increased in patients with preexisting atrial fibrillation

To investigate the potential mechanisms by which MRA reduces the occurrence of AF, we performed snRNA seq on left atrial tissue from organ donors without AF, patients with AF, and donor hearts that underwent cold preservation and perfusion with or without MRA (canrenone) treatment. After filtering the nuclei with low-quality nuclei, doublets, and ambient RNA contamination, we obtained 135,547 nuclei that were clustered into 10 distinct clusters. We annotated these clusters into Adipocytes, Cardiomyocytes, Endothelial cells, Fibroblasts, Macrophages, Mesothelial cells, Neuron, Smooth muscle cells, T cells and Pericytes based on the marker gene expression (Fig 3A-B and Fig S6A). The cellular composition in AF is associated with a proportional increase in mesothelial cells (Fig S6B, P = 0.026). Moreover, we found that MR (NR3C2) expression was significantly increased in atrial endothelial cells, fibroblasts, macrophages, and mesothelial cells. In human hearts that were cold static preserved for 10 hours followed by ex-vivo reperfusion, canrenone treatment significantly inhibited the global atrial expression of MR target genes (Fig 3D). MRA (canrenone) suppressed MR target genes were conversely increased in the left atrium of patients with chronic atrial fibrillation (Fig 3D).

**Figure 3.**
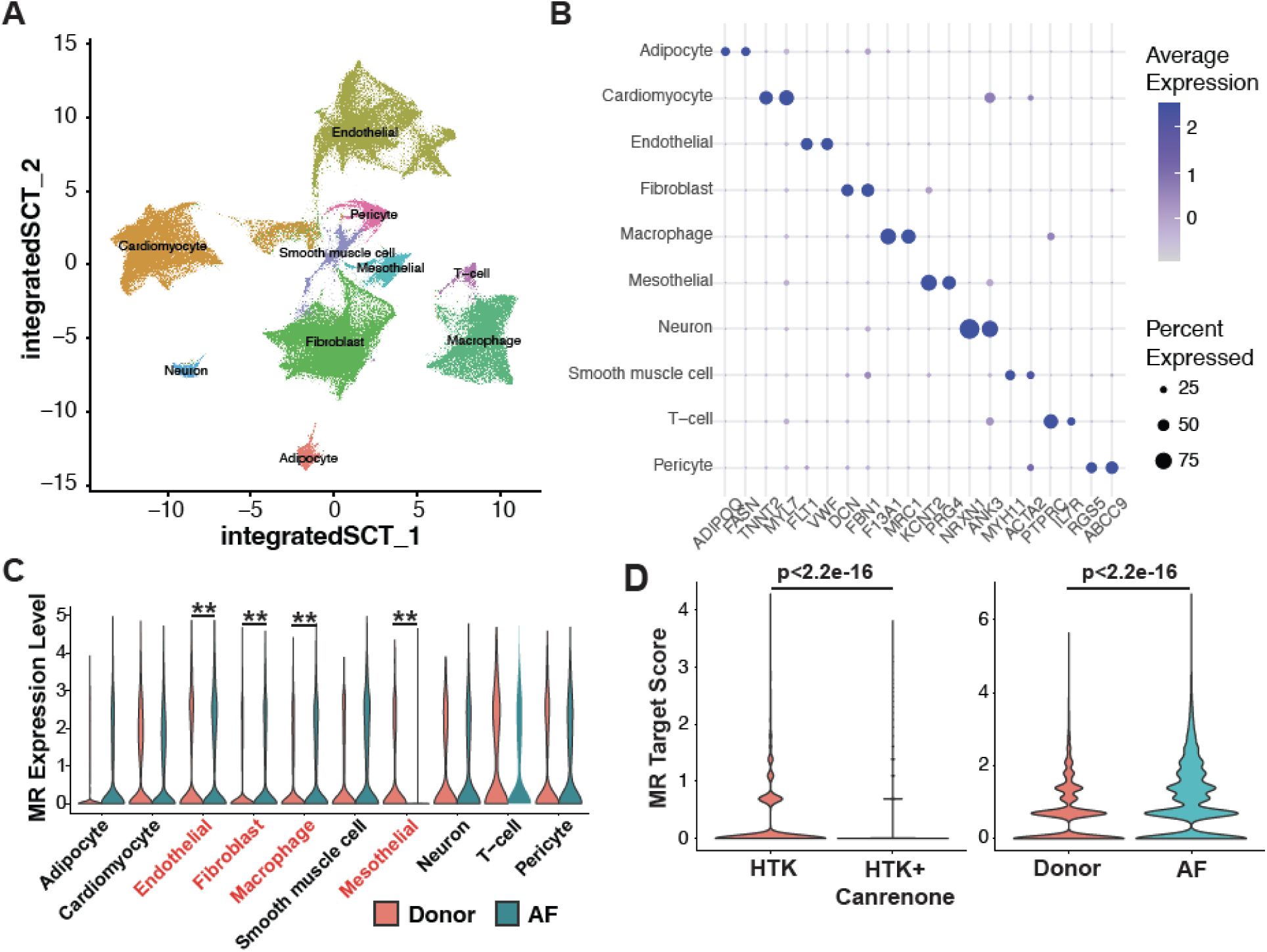
(A) Representative uniform manifold approximation and projection (UMAP) plot of snRNA-seq of LA tissues from different donors with integrated SCT reduction. 10 different cell types were clustered and identified. (B) Dot plots of the cell-specific marker gene expression. (C) MR (NR3C2) expression in each cell type in donor and AF patients was shown in the violin plot. ** and cell types in red letters indicate P < 0.001. (D) Violin plot of MR target expression in the left atrium of reperfused donor heart with or without canrenone treatment as well as donor and AF patients. CM – Cardiomyocytes.

### Canrenone inhibits activation of specific MR genes in human atrial cardiomyocytes which are associated with cold cardiac preservation and chronic atrial fibrillation

Given the key role of cardiomyocytes in electrical signal transduction^39^ and atrial fibrillation pathogenesis^40–42^, we performed recluster analysis on total CMs and identified 2 groups of cardiomyocytes (Fig 4A). AF patients had an increase in the CM2 population compared to non-AF donors (Fig. 4B). Interestingly, the CM2 population was enriched for gene expression related to inflammation and cellular stress such as “apoptosis”, “interferon_gamma_response” and “unfolded protein response” gene sets (Fig 4C). CM2 cardiomyocytes were also enriched for MR downstream genes (Fig 4D, P=0.0002). Importantly, addition of canrenone to histidine-tryptophan-ketoglutarate (HTK) preservation solution in preserved-reperfused human hearts reduced atrial cardiomyocytes MR target gene expression compared to HTK only control in the same MR geneset^43^ (Fig. 4E, P=0.027). Furthermore, canrenone suppressed cardiomyocyte expression of a gene subset normally induced by cold cardiac preservation. These genes are conversely elevated in patients with chronic AF (Fig. 4F). However, canrenone activated genes were also elevated in patients with chronic AF (Fig. 4G). These cardiomyocyte genes suppressed by canrenone may represent mechanisms by which MRAs can prevent POAF following cardiac preservation.

**Figure 4.**
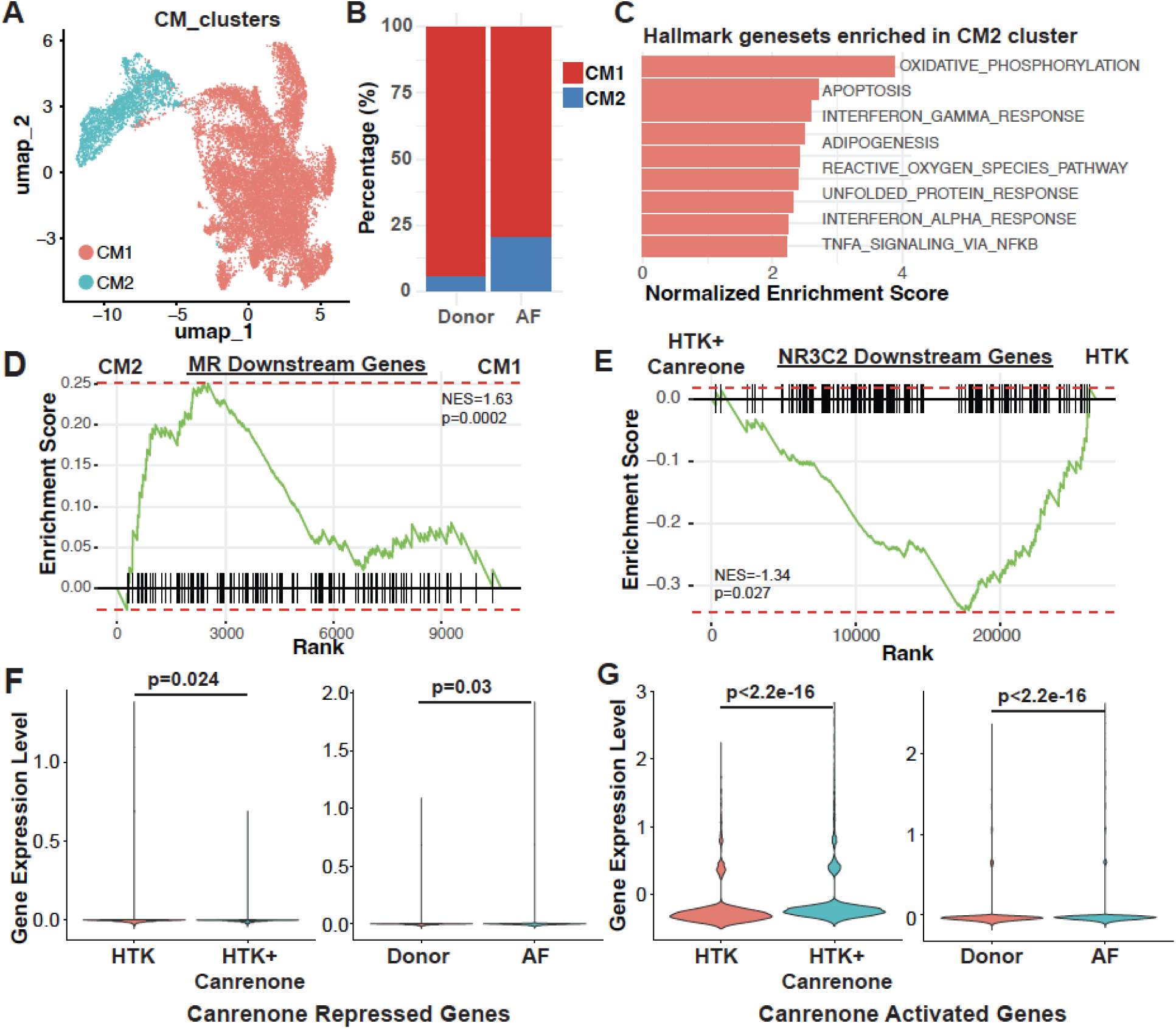
(A) Recluster of cardiomyocytes showed in UMAP plot. (B) Percentage of cardiomyocyte subclusters in donor vs AF patients. (C) GSEA analysis showed enriched gene sets in the CM2 compared to CM1. (D) GSEA of NR3C2 downstream gene set expression in CM2 compared to CM1. (E) GSEA showing effects from canrenone treatment of cold preserved-reperfused hearts on NR3C2 downstream genes in cardiomyocytes. (F) Violin plot of canrenone repressed expression of global gene expression in the LA of reperfused donor heart with or without canrenone treatment as well as donor and AF patients. (G) Violin plot of canrenone activated expression of global gene expression in the left atrium of reperfused donor heart with or without canrenone treatment as well as donor and AF patients. EC – Endothelial Cells.

### Canrenone suppresses human atrial macrophage specific gene activation associated with cardiac preservation and chronic atrial fibrillation

Given the key role of macrophages in chronic atrial fibrillation^44^, we also reclustered the macrophage population from combined donor and AF samples thus revealing 3 different subclusters (Fig 5A). There were no significant differences in the relative composition of macrophage subsets between the donor and AF atrial samples (Fig 5B). We then examined the differential expression genes between the preserved-reperfused donor hearts with and without canrenone. Interestingly, macrophage expression of genes that were repressed by canrenone are significantly higher in patients with chronic AF (Fig 5C). However, genes that were activated by canrenone following preservation-reperfusion were also activated in AF atria (Fig 5D). In the human preserved-reperfuesed hearts for example, canrenone suppressed macrophage associated FKBP5^45,46^ and ZBTB16^47,48^ expression which are elevated in the setting of chronic AF (Fig 5E). Both are known key genes associated with cellular stress, cell death, inflammation as well as AF. However, we did not see any difference in MR geneset expression in AF versus donor atria (Fig. 5F) nor in cold preserved-reperfused donor heart with or without canrenone treatment (Fig. 5G).

**Figure 5.**
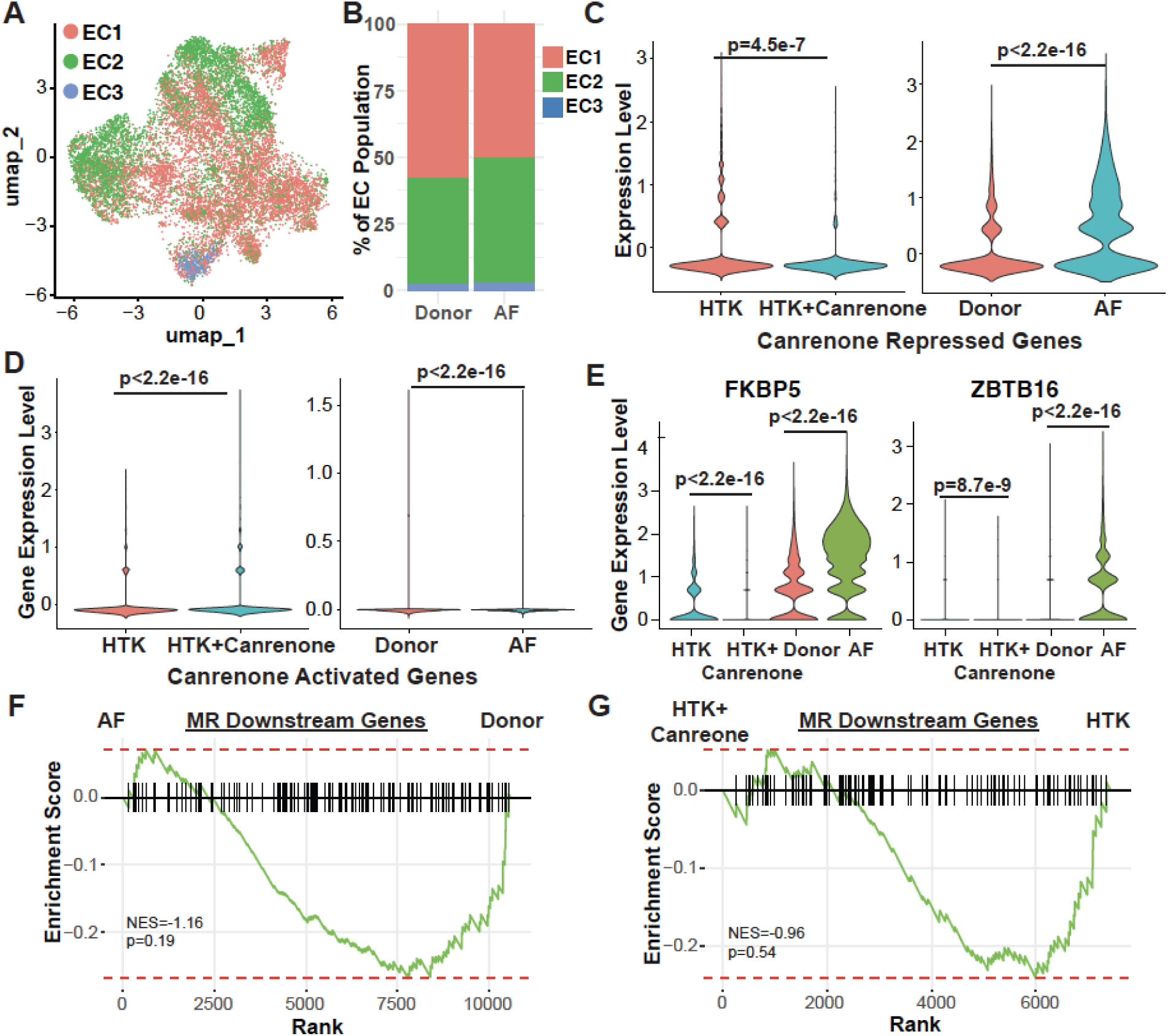
(A) Recluster of macrophage showed in UMAP plot. (B) Percentage of macrophage subclusters in donor vs AF patients. Violin plot of canrenone (C) repressed and (D) activated genes in macrophage of reperfused donor atria with or without canrenone treatment as well as in atria of donor heart and AF patients. (D) Violin plot shows FKBP5 and ZBTB16 expression. (D) GSEA showing macrophage expression of NR3C2 downstream gene set in AF versus donor atria. (E) GSEA showing the effects of canrenone treatment on NR3C2 downstream gene expression in macrophages.

### Vascular pericytes, but not endothelial and smooth muscle cells, show canrenone-induced suppression of genes that are associated with atrial fibrillation

Atrial endothelial cells (ECs) from combined donor and AF atrial samples were reclustered into 8 different subsets (Fig S7A). The atrial EC1 population was significantly enriched in AF patients (Fig S7B,). EC1 cells had enriched gene sets related to adipogenesis, and interferon alpha/gamma responses (Fig S7C). EC expression of canrenone-repressed (Fig. S7D) gene were increased in atria of AF patients. However canrenone-activated genes (Fig. S7E) were also activated in chronic AF atria. There was no significant enrichment of MR signals in the EC1 population compared to other EC subclusters (Fig S7F, P=0.38). The addition of canrenone did not impact endothelial expression of MR gene targets compared to HTK-only treated hearts (Fig. S7G, P=0.81).

In atrial smooth muscle cells, we identified MRA (canrenone) repressed genes in smooth muscle cells that were also similarly decreased in AF patients compared to donor atria (Fig. S8A). Similarly, MRA (canrenone) activated genes in smooth muscle cells were also increased in atria of AF patients compared with donors (Fig. S8B). Smooth muscle cell expression of MR target genes were also not different between atria of AF patients (Fig. S8C) and donors nor did canrenone treatment impact their expression in cold preserved-reperfused donor hearts (Fig. S8D).

In atrial pericytes, MRA (canrenone) repressed and activated genes that are conversely up (P=0.036) and downregulated (P=0.00067) in atrial pericytes from AF patients (Fig. S8E-F). While atria of AF patients had higher expression of MR target genes compared to donors (Fig. S8G), canrenone treatment did not impact pericyte expression of MR target genes in cold preserved-reperfused hearts. Findings from the vascular cell populations suggest that the EC and smooth muscle cells may not be the target of canrenone for preventing POAF. However, MRA treatment of cold preserved hearts did impact pericyte expression of genes that conversely demonstrate mirrored expression in the atria of patients with AF.

## DISCUSSION

While MRAs are known to reduce the development and burden of AF in chronic medical settings^15–19^, initiating MRA in the acute setting for reducing POAF is less clear. Using propensity matching to obtain more similar comparison patients populations, we show that preoperative MRA use in the total propsensity matched population reduced POAF from 31.5% to 19.8%. Even for propensity matched populations who used preoperative diuretics, POAF was reduced from 33.2% to 19.8% suggesting that MRA has effects on reducing POAF that is independent of volume reduction. Importantly, patients with POAF are more prone to develop chronic atrial fibrillation long term. Bianco et al reported that the POAF cohort had a significantly higher incidence of atrial fibrillation on follow-up for up to 5 years (11.74% vs 4.75%; *P* <0.001)^49^. In the current study, preoperative MRA use was associated with a 40.7% reduction in the development of paroxysmal or chronic AF at 6 years. This suggests that preoperative MRA use should be further studied as a strategy to not only reduce POAF short term but also reduce the longer term morbidity of chronic AF.

MR activation as a mechanism for POAF have been implicated in the ALDO-POAF study (ALDOsterone for prediction of Post-Operative Atrial Fibrillation, NCT 02814903) where higher preoperative aldosterone levels was shown to be a biomarker for identifying patients susceptible to developing POAF after ccoronary artery bypass grafting (CABG) with or without aoric valve replacement (AVR)^50^. This implies a key pathogenic role of MR activation in POAF^50,51^. Based on this rationale, Pretorius et al^52^ randomized 445 patients undergoing randomization to preoperative oral ramipril (2.5 – 5 mg daily), spironolactone (25 mg daily) or placebo for 4-7 days prior to elective CABG or valve surgery. Although preoperative MRA was associated with less postoperative renal failure and quicker extubation, there was unfortunately no difference in POAF rates between the groups^52^.

However, limitations in the trial by Pretorius et al^52^ include being inadequately powered since it recruited 147 to 151 patients per group for a relatively low incidence of POAF of 27% in their study population. Furthermore, this study included patients who had paroxysmal or chronic AF as long as it did not occur within 6 months of the enrollment. It is also likely that in this baseline MRA naïve population, there is a failure to reach therapeutic circulating MRA threshold levels for inhibtion of cardiac inflammation and oxidative stress during cardioplegic arrest. Our unpublished studies show that improved cardiac preservation quality requires a intracardiac dosing of canrenone, an MRA metabolite of spironlactone, at a concentration of 50 μM given along with cardioplegia during cardiac surgery. The ATHENA-HF trial for determining the efficacy of oral MRA (spironolactone) for treating acute heart failure gave important insight into challenges of oral spironolactone treatment duration and dose for reaching therapeutic systemic level of spirononlactone active metabolites. It was shown that even at spironolactone 100 mg oral daily, many of its active metabolites (including canrenone) remained at very low circulating concentrations after 96 hours in MRA naïve patients^53^. Patients who were on spironolactone previously had significantly higher concentrations^53^ suggesting that oral spironolactone administration would take much longer than 1 week to reach therapeutic steady state concentration at a lower 25 mg dosage in MRA naïve patients in the study by Pretorius et al^52^. Direct intracardiac MRA (canrenone) administration by addition ot cardioplegic solution during cardiac surgery is expected to overcome the limitations of patient compliance and achieving therapeutic MRA levels in cardiac tissue.

To further characterize the effects of MRA on gene expression in the left atrial cell population in the setting of cold cardioplegic arrest and cardiac preservation during cardiac surgery, we performed single-nuclei RNA sequencing (snRNA-Seq) analysis. Using canrenone, a water soluble MRA, we determine the cell populations that affected by MR inhibition in terms of AF associated gene expression obtained by comparing gene expression between the atria of patients with AF and donors without AF. Canrenone, a water-soluble active metabolite of spironolactone, was able to suppress the upregulation of MR target genes in cold preserved donor hearts and these MR genesets are conversely upregulated in the left atrium of patients with chronic AF. This correlates with our clinical finding that preoperative MRA is able to reduce POAF incidence and long term AF burden.

Cardiomyocytes are known to serve an important role in cardiac electrical signal transduction^39^, and atrial fibrillation pathogenesis, including in the postoperative setting^40–42^. Therefore, we focused on the human atrial cardiomyocyte population showing the human CM2 cardiomyocytes are enriched for MR expression consistent with their preferred activation of signaling pathways relevant for inflammation, cellular stress and cell death. Canrenone was able to suppress preservation associated MR activation within atrial cardiomyocytes suggesting intraoperative canrenone added to cardioplegic solution used for cardiac preservation during cardiac surgery may reduce POAF burden. As indicated in our global gene expression analysis in cardiomocytes, canrenone is also able to reduce the expression of other non-MR target genes that may be relevant for AF pathogenesis.

Recent reports have also indicated macrophages as playing a key role in AF^44^. Although there was no difference in the proportion of macrophage subsets, cardiac preservation using canrenone did suppress macrophage expression of genes that are elevated in atrial macrophages of patients with chronic AF. Examples include suppression of FKBP5 and ZBTB16 which have key role in cellular survival, stress responses, inflammation as well as specially in AF pathogenesis. However, we did not detect an impact by canrenone on MR geneset expression in macrophages. Canrenone also suppressed expression of a subset of cell specific genes in endothelial smooth muscle cells and pericytes, that were elevated in the respective cells in chronic AF. However, canrenone activated genes inthese vascular cell types were also activated in the atria of patients with chroic AF. This suggests that cardiomyocytes are likely key targets for canrenone suppression of AF through MR antagonism. However, However, it is possible that canrenone may also act on macrophages and vascular cells to reduce AF although this is more likely through a MR independent or nongenomic mechanism^54^ since it did not impact MR geneset expression.

### Study limitations

Our study remains limited by its single institution retrospective design with associated biases despite use of propensity matching to obtain more comparable groups. Furthermore, POAF may also occur after patient discharge but are not be detected due to a lack of symptoms or patients not presenting for evaluation will be missing from our dataset. It is also possible that the effect of MRA reducing the incidence of POAF results from chronic effects of antagonizing structural remodeling^21,55,56^ leading to a less dilated atrium and/or attuenating electrical remodeling of atrial tissue^21,55^ rather than inhibiting the acute effects of cold preservation on the atria. Moreover, a systemic proinflammatory milieu from the effects of cardiopulmonary bypass^57,58^ and/or general physiological stress from surgical trauma^58^ may impact propensity for POAF independent of hyperactivation of MR within the heart from cold preservation injury. Indeed, the sources of stimulus for POAF in the cardiac surgery setting likely differs from that of nonsurgical paroxysmal or chronic AF. Our snRNA-seq data are suggestive of the role of MR signaling in POAF by correlating with gene expression signature of patient with chronic AF. We have not taken atrial tissue from patients who experience new POAF versus those who did not since atrial tissue is generally not removed unless there is pre-existing AF. Removal of the left atrial appendage in the setting of pre-existing AF is done to reduce stroke risk^59^.

## Conclusions

We present propensity matched retrospective data that show an important association between preoperative use of oral MRA with significantly reduced POAF and longer term development of persistent AF after cardiac surgery utilizing cardiopulmonary bypass. Furthermore, single-nuclei RNA sequencing studies suggests that canrenone added to cardiac preservation solution can inhibit signaling pahways related to chronic AF including MR activation. Future mechanistics studies leveraging genetic MR deficiency or knockdown in an AF prone model to specifically determine the role of MR activation in POAF. Furthermore, these findings support the conduct of future clinical trials to determine the role of preoperative MRA therapy and/or canrenone added to cardioplegia solution with the goal of reducing POAF incidence as well as subsequent development of AF long term.

## Acknowledgements

The authors thank organ donors and their families, patients undergoing cardiac surgery, as well as organ procurment organizations (Gift of Life Michigan and LifeSource of Minnestoa) for the opportunity to obtain human atrial samples for this study. We acknowledge the Mayo Clinic for supporting this study. We also thank Jolene Erola, Erika Trower and Becky Wait for administrative support. We also greatly appreciate the Mayo Clinic Department of Pathology, including Monica Kendall, P.A., Luke Wilson, P.A., and Joseph J. Maleszewski, M.D., for tissue procurement and processing.

## Abbreviations List

AF: Atrial fibrillation
EC: Endothelial cell
HTK: Histidine-Tryptophan-Ketoglutarate
MR: Mineralocorticoid receptor
MRA: Mineralocorticoid receptor antagonist
POAF: Postoperative atrial fibrillation
STS: Society of Thoracic Surgeons
STS-ACSD: Society of Thoracic Surgeons Adult Cardiac Surgery Database

